# Intraclass correlation: improved modeling approaches and applications for neuroimaging

**DOI:** 10.1101/164327

**Authors:** Gang Chen, Paul A. Taylor, Simone P. Haller, Katharina Kircanski, Joel Stoddard, Daniel S. Pine, Ellen Leibenluft, Melissa A. Brotman, Robert W. Cox

## Abstract

Intraclass correlation (ICC) is a reliability metric that gauges similarity when, for example, entities are measured under similar, or even the same, well-controlled conditions, which in MRI applications include runs/sessions, twins, parent/child, scanners, sites, etc. The popular definitions and interpretations of ICC are usually framed statistically under the conventional ANOVA platform. Here, we provide a comprehensive overview of ICC analysis in its prior usage in neuroimaging, and we show that the standard ANOVA framework is often limited, rigid, and inflexible in modeling capabilities. These intrinsic limitations motivate several improvements. Specifically, we start with the conventional ICC model under the ANOVA platform, and extend it along two dimensions: first, fixing the failure in ICC estimation when negative values occur under degenerative circumstance, and second, incorporating precision information of effect estimates into the ICC model. These endeavors lead to four modeling strategies: linear mixed-effects (LME), regularized mixed-effects (RME), multilevel mixed-effects (MME), and regularized multilevel mixed-effects (RMME). Compared to ANOVA, each of these four models directly provides estimates for fixed effects as well as their statistical significances, in addition to the ICC estimate. These new modeling approaches can also accommodate missing data as well as fixed effects for confounding variables. More importantly, we show that the MME and RMME approaches offer more accurate characterization and decomposition among the variance components, leading to more robust ICC computation. Based on these theoretical considerations and model performance comparisons with a real experimental dataset, we offer the following general-purpose recommendations. First, ICC estimation through MME or RMME is preferable when precision information (i.e., weights that more accurately allocate the variances in the data) is available for the effect estimate; when precision information is unavailable, ICC estimation through LME or the RME is the preferred option. Second, even though the absolute agreement version, ICC(2,1), is presently more popular in the field, the consistency version, ICC(3,1), is a practical and informative choice for whole-brain ICC analysis that achieves a well-balanced compromise when all potential fixed effects are accounted for. Third, approaches for clear, meaningful, and useful result reporting in ICC analysis are discussed. All models, ICC formulations, and related statistical testing methods have been implemented in an open source program `3dICC`, which is publicly available as part of the AFNI suite. Even though our work here focuses on the whole brain level, the modeling strategy and recommendations can be equivalently applied to other situations such as voxel, region, and network levels.

## Introduction

Recently, reliability and reproducibility have been hot topics in science in general and in the neuroimaging community in particular. It is known that neuroimaging data are very noisy and a large proportion of the FMRI variability cannot be properly accounted for: It has been reported that less than half of the data variability can currently be explained in the typical data analysis (Gonzalez-Castillo et al., 2017). For example, the typical analytical approach is to make a strong and unrealistic assumption that a hemodynamic response is the same across brain regions, subjects, groups, different tasks or conditions. In addition, even though large amounts of physiological confounding effects are embedded in the data, it remains a daunting task to fully incorporate the physiological noise in the model.

Recent surveys observed that about 60% of published experiments failed to survive replication in psychology (Baker, 2015) and about 40% in economics (Bohannon, 2016), and the situation with neuroimaging is likely to be equally, if not more, pessimistic (Griffanti et al., 2016). While in neuroimaging, reproducibility is typically a qualitative description as to whether an activation cluster is reported across many studies, the reliability of an effect estimate quantitatively describes the variation in repeated measurements performed on the same measuring entities (e.g., subjects in neuroimaging) under the identical or approximately the same experimental conditions (NIST, 2007). Specifically, reliability can be defined as the agreement or consistency across two or more measurements, and intra-class correlation (ICC) has been specifically developed for this purpose (Shrout and Fleiss, 1979; McGraw and Wong, 1996).

Generally speaking, the conventional ICC metric indicates agreement, similarity, stability, consistency, or reliability among multiple measurements of a quantity. For instance, such a quantity can be the ratings of *n* targets assessed by *k* raters or judges in a classical example (Shrout and Fleiss, 1979). In the neuroimaging context, when the same set of measuring entities (e.g., subjects) go through the same experiment protocol under the same conditions, ICC can be utilized to assess the data quality. Those multiple measurements can be the effect estimates from *n* subjects under *k* different replications (e.g., runs, sessions, scanners, sites, twins, siblings, parent-child pairs, studies, assessments, diagnoses, or analytical methods). The same quantity (rating or effect estimate) across those *k* replications in the ICC definition is reflected in the word *intra*class, as opposed to the Pearson (or *inter*class) correlation coefficient that reveals the linear relationship between two quantities that can be of different nature (e.g., brain response and cortex thickness).

Here we first review the various versions of ICC definition and their computation formulations under their classic ANOVA platform, and then we discuss their limitations and drawbacks as our motivations to develop more extended models. We then describe and validate several new, improved approaches for ICC estimation. Even though our work mainly focuses on whole brain data analysis in neuroimaging, the methodologies can be applied to other contexts or fields when the underlying assumptions are met.

### Various types of ICC

In this section, we introduce the three classic types of ICC, motivating each from their basic statistical model and describing their interpretation, applicability, and generalization. Throughout this article, regular italic letters in lower case (e.g., a) stand for scalars and random variables; boldfaced italic letters in lower (***a***) and upper (***X***) cases for column vectors and matrices, respectively; Roman and Greek letters for fixed and random effects, respectively, on the righthand side of a model equation. In the neuroimaging context, let *y*_*ij*_ be the effect estimate at the ith level of within-subject (or repeated-measures) factor A and the jth level (usually subject) of factor B (*i* = 1, 2,…,*k*; *j* = 1, 2,… *n*). When both factors A (e.g., runs, sessions, scanners, sites) and B are modeled as random effects, we have a two-way random-effects ANOVA system,

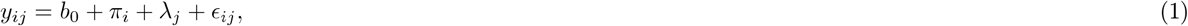

where *b_0_* is a fixed effect or constant representing the overall average, *π*_*i*_ is the random effect associated with the *i*th level of factor A, *λ*_*j*_ represents the subject-specific random effect, and ϵ*_ij_* is the residual. With the random variables *π*_*i*_, *λ*_*j*_, and *ϵ*_*ij*_ assumed to be independent and identically distributed with 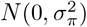, 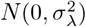, and 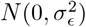, respectively, the associated ICC for the model (equation 1) is defined as

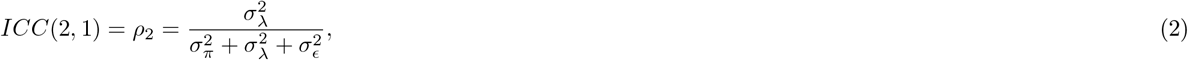

and is numerically evaluated by

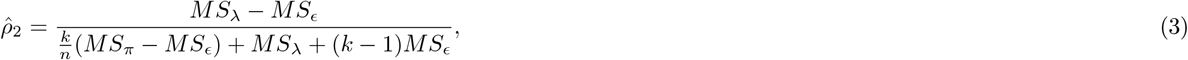

where *M S*_*π*_, *M S*_*λ*_ and *M S*_*ϵ*_ are the mean squares (*M S*s) associated with the factor A effects *π*_*i*_, subject effects *λ*_*j*_ and the residuals *ϵ*_*ij*_, respectively, in the ANOVA framework (equation 1). The definition (equation 3) is usually referred to as ICC(2,1) in the literature (e.g., Shrout and Fleiss, 1979; McGraw and Wong, 1996).

To make statistical inference, Fisher’s transformation for ICC value *ρ* (McGraw and Wong, 1996),

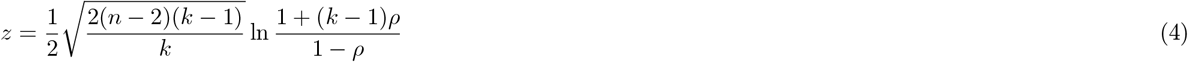

approximately follows a standard Gaussian *N*(0,1) under the null hypothesis *H*_0_: *ρ* =0, and offers a solution for significance testing. However, a better approach is to formulate an *F*-statistic (McGraw and Wong, 1996),

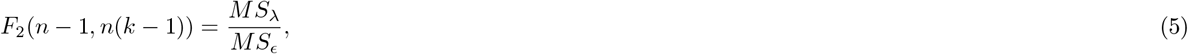

whose distribution is exact under *H*_0_: *ρ*_2_ = 0, unlike the Fisher transformation (4).

The meaning of the ICC can be interpreted in four common perspectives:

i. As the definition (2) itself indicates, the ICC is the proportion of total variance that is attributed to a random factor (or accounted for by the association across the levels of the random factor). For instance, if the variance associated with subjects increases, subjects would be less similar while the levels of factor A (e.g., runs) tend to be relatively more similar, leading a higher ICC value. This proportionality interpretation is straightforwardly consistent with the non-negativity of ICC and its range of [0,1].
ii. The ICC is the expected correlation between two effect estimates that are randomly drawn among the levels of factor A within the same level of factor B. For instance, say that ICC(2,1) for an FMRI study shows the relatedness among multiple runs. Specifically, with the assumptions in the model (1), ICC(2,1) is essentially the Pearson correlation of the effect estimates between any two levels (e.g., runs) of factor A, *i*_1_ and *i*_2_ (*i*_1_ ≠ *i*_2_),

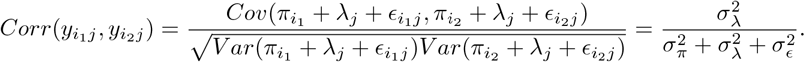 However, it is worth emphasizing that this equivalence between ICC and Pearson correlation holds because of the following fact: ICC is a relationship gauge between, for example, any two runs in light of the same physical measure (e.g., BOLD response in ICC(2,1) as opposed to Pearson correlation between, for example, weight and height). When a run has generally a higher (or lower) effect estimate relative to the group mean effect, or when there is some extent of consistency among subjects within each run, then those effect estimates are correlated, and the ICC formulation (2) basically captures that correlation or consistency.
iii. ICC is an indicator of homogeneity of the responses among the levels (e.g., subjects) of the random factor B: a higher ICC means more similar or homogeneous responses. On the other hand, when an ICC value is close to zero, the effect estimates associated with a factor A level are no more similar than those from different subjects, and the random effect components could be removed from the model (1).
iv. ICC reflects the extent of common conditions (e.g., same task and scanning parameters) that the effect estimates share. The ICC would be higher if effect estimates associated with a subject were under more similar environments.

The decision of an explanatory variable in a model as either fixed or random effects can be subtle, and the distinction is usually determined in light of the nature of the factor: interchangeability of factor levels, or whether there exists some systematic difference across the factor levels. For example, subjects are often considered as the levels of a random factor because: 1) they are each recruited through a random sampling process as representatives (or samples) of a potential or conceptual population (as embodied in the assumption of Gaussian distribution 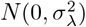 for the random effects *λ*_*j*_ in the model (1)), achieving the goal of generalization in statistical inferences; 2) their order does not matter in the model and can be randomly permuted due to exchangeability; and 3) a particular set of subjects can, in practice, be replaced by another set. In contrast, patients and controls are typically handled as fixed effects because of the lack of exchangeability. In neuroimaging scanners or sites can be thought of as the levels of either a random- or fixed-effects factor, depending on whether the scanning parameters are similar or different across scanners or sites. Similarly, runs or sessions should be treated as random effects if no significant systematic difference exists across runs or sessions; otherwise, they should be modeled as fixed effects when habituation or familiarity effect of the task is substantial.

Because of the distinction between fixed and random effects, there is an alternative ICC definition in which the factor A (e.g., runs, sessions, scanners, sites) is modeled as fixed effects in a two-way mixed-effects ANOVA structure,

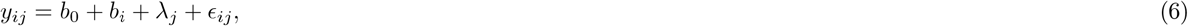

where *b*_*i*_ and *λ*_*j*_ represent the fixed effects of factor A and random effects of subjects (or families, in the case of parent versus child), respectively. The associated ICC is defined as

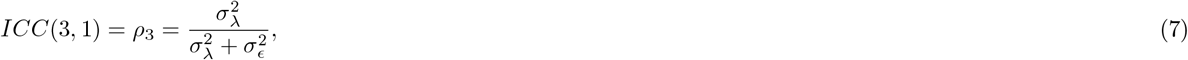

which has the same formula as (2), but we note that the presence of *b*_*i*_ here means that the two models would have different estimates of 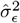 The ICC in this case can be computed as

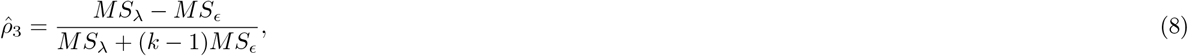

with an exact *F*-statistic,

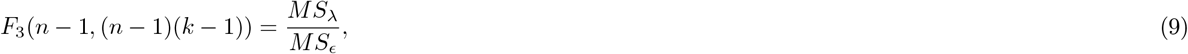

for significance testing under the null hypothesis *H*_0_: *ρ*_3_ = 0.

The two ICC definitions, (2) and (7), are popularly notated as ICC(2,1) and ICC(3,1), respectively, and are sometimes referred to as “intertest ICCs,” extensions of the classic inter-rater reliability (Shrout and Fleiss, 1979). When there are only two levels for factor A (*k* = 2), these two versions are usually referred to as test-retest reliability measures. Yet there is another ICC type in which the effects of factor A are not explicitly modeled. A prototypical example is the scenario with the effect estimates of twins from each family where there is no meaningful way to assign an order or sequence among the levels of factor A consistently among the levels of factor B (e.g., ordering twins within each family). Let *y*_*ij*_ be the effect estimate (BOLD response in percent signal change or connectivity measurement) from the *i*th level of factor A and *j*th level of factor B (e.g., the *i*th member of family *j*)(*i* = 1,2,…,*k*;*j* = 1,2,…,*n*). A one-way random-effects ANOVA can be formulated to decompose the response variable or effect estimate *y*_*ij*_ as

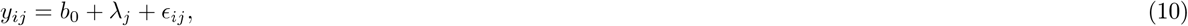

where *λ*_*j*_ codes the random effect of the *j*th level of factor B (e.g., family *j*).

The ICC for the model in (10) is defined as

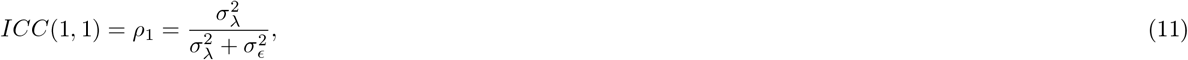

which can be estimated similarly as (8),

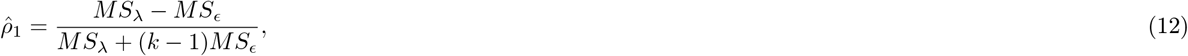

where *M S*_λ_ and *M S*_*ϵ*_ are the mean squares (*M S*s) associated with the family effects *λ*_*j*_ and the residuals, respectively, in the ANOVA framework of model (10). The testing statistic for (11) has the form as (5), and the definition (11) is usually referred to as ICC(1,1) in the literature (e.g., Shrout and Fleiss, 1979; McGraw and Wong, 1996).

The four interpretation perspectives for ICC(2,1) also apply directly to other two types. For example, with the assumptions in the model (6), ICC(3,1) is a special case of Pearson correlation of the effect estimates between any two levels of factor A, *i*_1_ and *i*_2_ (*i*_1_ ≠ *i*_2_),

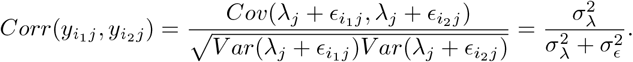

Nevertheless, ICC(3,1) is more similar to Pearson correlation than ICC(1,1) and ICC(2,1) in the sense that each level of factor A is assumed to have a different mean, but it remains unlike Pearson correlation as the assumption of same variance holds across the levels of factor A for all the three ICC definitions. In addition, ICC(2,1) and ICC(3,1) are sometimes called *absolute* agreement and *consistent* agreement, respectively. The distinction between these two ICC types can be hypothetically illustrated by paired effect estimates (e.g., in percent signal change for an FMRI experiment) from five subjects during two sessions (Table 1). The three ICC values are all nonnegative because they represent a proportion of total variance embedded in the data, and they generally follow a sequential order (Shrout and Fleiss, 1979): *ICC*(1,1) ≤ *ICC*(2,1) ≤ *ICC*(3,1).

**Table 1:** Hypothetical effect estimates (e.g., BOLD response in percent signal change) with a relationship from five subjects during two sessions (*y*_2*j*_ = *y*_1*j*_ + 0.2,*j* = 1, 2,…,5; ICC(1,1) = 0.43, ICC(2,1) = 0.56, ICC(3,1) = 1, Pearson correlation *r* = 1).

| subject | s1 | s2 | s3 | s4 | s5 |
| --- | --- | --- | --- | --- | --- |
| session 1, $y_{1j}$ | 0.1 | 0.2 | 0.3 | 0.4 | 0.5 |
| session 2, $y_{2j}$ | 0.3 | 0.4 | 0.5 | 0.6 | 0.7 |

The first index in the ICC(·,·) notation specifies the ICC type, while the second indicates the relationship between two *single* measurements (e.g., between two twins for ICC(1,1) or two levels of factor A for ICC(2,1) and ICC(3,1)). For each of the three *single measurement* ICC types, there is another version, called *average measurement* ICC, which shows the relationship between two sets of average measurements among the *k* levels of factor A, and with notations ICC(1,*k*), ICC(2,*k*), and ICC(3,*k*), they are similarly defined as the single measurements version except that the terms in the denominator,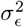,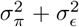 and 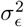, for ICC(1,1), ICC(2,1) and ICC(3,1), respectively, are each scaled by a factor of *k*^−1^. By definition, the average measurements ICC is larger than its single measurements counterpart. In addition, a similar correlation interpretation about the average measurement ICCs can be seen with, for example, ICC(2,*k*),

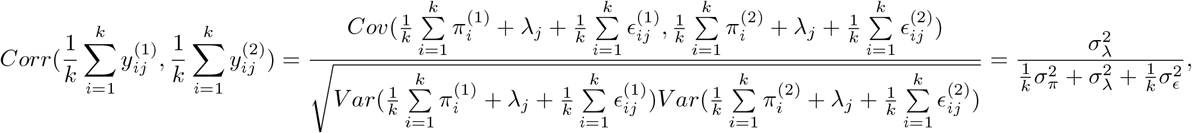

where the superscript such as those in 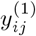 and 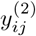 indicates a particular set of data substantiation, and thus 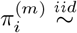,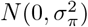,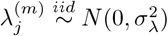,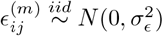. As the average measurement ICC is less popular in practice, we hereafter focus on the their single measurement counterpart, but our modeling work below can be directly extended to the average measurements ICC.

### Literature survey of ICC for neuroimaging

ICC has been applied to neuroimaging data for over 10 years, mainly to examine reliability under various scenarios. In particular, ICC(2,1) has been largely adopted in the field, using the ANOVA approach. ICC(2,1) has been used to show reliability under various scenarios, for example, at the regional level (Fiecas et al., 2013), at the network level for resting-state (Guo et al., 2012; Cao et al., 2014), at the whole brain level with ANOVA (Kristo et al., 2014; Zanto et al., 2014; Quiton et al., 2014) and with linear mixed-effects (LME) modeling using a precursor of `3dICC` in AFNI (Fiecas et al., 2013; White et al., 2016; Haller et al., 2017). ICC(3,1) has been applied at the regional level for task-related FMRI data (Cáceres et al., 2009; Jaeger et al., 2015) and for MEG data (Recasens and Uhlhaas, 2017), at the network level for resting-state data (Braun et al., 2012), and at the whole brain level for task-related FMRI data, using ANOVA in PASW and/or Matlab (Cáceres et al., 2009; Plichta et al., 2012; Brandt et al., 2013) and in SAS (Fournier et al., 2014). It should be noted that in many cases the ICC type adopted in a study was not clearly stated, often due to the ambiguity in terms of the model involved (Zuo et al., 2010b; Lin et al., 2015; Shah et al., 2016).

There have been occasions in which ICC(1,1) and ICC(3,1) have been explicitly employed at times in the literature. For example, ICC(1,1) has been applied to brain networks based on resting-state FMRI data (Wang et al., 2011), to functional near-infrared spectroscopy (fNIRS) data (e.g., Bhambhani et al., 2006; Pichta et al., 2006; Pichta et al., 2007; Zhang et al., 2011; Tian et al., 2012), and to resting-state data at the whole brain level (Zuo et al., 2010a), as well as to task-related data through LME at the regional level (Töger et al., 2017). However, we note that in a large number of publications, the adoption of the ICC type was neither explicitly explained nor justified, which makes precise interpretation difficult.

On the whole brain level, in addition to the ICC computation through LME as implemented in the open source AFNI program `3dLME` (Chen et al., 2013), there have been a few Matlab toolboxes publicly available: three using ANOVA (Cáceres et al., 2009; Fiecas et al., 2013; Molloy and Birn, 2014), and one using both ANOVA and LME in Matlab (Zuo et al., 2010b). The concept of ICC has also been extended to characterize cross-subject heterogeneity when the effect estimate precision is incorporated into FMRI group analysis (Chen et al., 2012), as well as to reveal the relatedness among subjects in inter-subject correlation (ISC) analysis with data from naturalistic scanning (Chen et al., 2017a).

### Motivations for further modeling work

The ANOVA framework has been adopted in computing ICC through the MS terms largely for historical reasons because ANOVA was developed and became widely adopted in early 20th century: the framework is widely introduced in basic statistics textbooks, and the MS terms are efficient to compute. Various computational tools are widely available through ANOVA (e.g., packages *irr* and *psych* in R). However, the ANOVA approach does have both limitations for interpretation and practical drawbacks for calculation:

i. *Negative value.* Although the ICC value should be theoretically nonnegative per its definition as the proportionality of the total variance (as in (2)), its estimation (as in (3)), may become negative due to the fact that the numerator in the computational formula is the difference between two MS terms. Such cases are uninterpretable. Importantly, in neuroimaging negative ICCs are not rare occurrences, with a large number of such voxels appearing in the brain (and in any tissue type).
ii. *Missing data.* In common practical situations, missing data may occur. As data balance is essential in partitioning the MS terms, ANOVA cannot properly handle missing data due to the breakdown of the underlying rigid variance-covariance structure. For example, the data for the six subjects who missed scanning for one session as shown in Table 2 would have to be abandoned due to the rigidity of the ANOVA structure.

**Table 2.** A hypothetical study of 17 subjects with missing data (1: available, 0: missing)
iii. *Confounding effects.* Under some circumstances it might be desirable to incorporate explanatory variables into a model, so that the variability due to those confounding effects can be properly accounted for. For example, subject groupings (e.g., sex, handedness, genotypes), age, and reaction time are typical candidates in neuroimaging that often warrant their inclusion in the model. However, the rigid ANOVA structure usually does not easily allow for such inclusions.
iv. *Sampling errors.* Conceptually, the residual term *ϵ*_ij_ in the ICC model can be considered to represent measurement or sampling errors. In other words, the underlying assumption is that all the effect estimates *y*_*ij*_ share the same sampling variance for the measurement errors. However, unlike the typical situation in other fields where the effect estimates are usually direct measurements (e.g., scores assessed by raters), the effect estimates in neuroimaging are generally obtained through a data reduction process with a regression model for each subject. That is, each effect estimate is associated with a standard error that indicates the precision information of the estimation, and heterogeneity or (possibly unfounded) heteroscedasticity is expected because the standard error varies across subjects as well as between twins or parents/children, across runs/sessions or scanners/sites. When the standard error for the effect estimate is ignored in favor of a homoscedasticity assumption in the ANOVA formulation (as widely practiced in neuroimaging when computing the ICCs, as well as in group analysis), it raises the question: what is the impact for the ICC estimate when heterogeneity of standard error is substantially present across the measuring entities?
v. *Type selection.* The applicability of ICC(1,1) is limited to situations where the levels of the repeated-measures factor are measuring entities that are difficult to assign meaningful orders or sequences such as twins. However, between ICC(2,1) and ICC(3,1), the choice becomes challenging for the investigator, with nontrivial impact on the ICC results: other than some prior knowledge about the potential existence of confounding effects or systematic difference across the levels of the repeated-measures factor, there is no solid statistical tool under ANOVA to leverage one choice over the other. For example, is there any statistical evidence that could allow us to decide unequivocally between ICC(2,1) and ICC(3,1) by treating runs or sessions as random or fixed effects? In addition, the typical whole brain analysis through a massively univariate approach at the voxel level may further aggravate the choice.

| subject | s1 | s2 | s3 | s4 | s5 | s6 | s7 | s8 | s9 | s10 | s11 | s12 | s13 | s14 | s15 | s16 | s17 |
| --- | --- | --- | --- | --- | --- | --- | --- | --- | --- | --- | --- | --- | --- | --- | --- | --- | --- |
| session 1 | 1 | 1 | 1 | 1 | 1 | 1 | 1 | 1 | 1 | 1 | 1 | 1 | 1 | 1 | 1 | 1 | 1 |
| session 2 | 1 | 1 | 1 | 1 | 0 | 1 | 1 | 0 | 1 | 1 | 1 | 1 | 1 | 1 | 1 | 1 | 1 |
| session 3 | 1 | 0 | 1 | 1 | 1 | 0 | 1 | 1 | 1 | 1 | 1 | 0 | 1 | 1 | 0 | 1 | 1 |

Our ICC modeling work here hinges around these five limitations of the ANOVA approach. We first discuss four alternative modeling approaches as extensions to the ANOVA framework, and then use an experimental dataset to examine the performance of the various modeling methods. A literature survey and further discussion are presented at the end. The implementations of our modeling work are publicly available for voxel-wise computation through program `3dICC` as part of the AFNI suite (Cox, 1996). As ICC(1,1) is typically adopted in neuroimaging for special cases of studies with twins, our focus here is on the other two types due their wider applicability. Nevertheless, the modeling strategies discussed here can be directly expanded to ICC(1,1).

## Theory: three extended ICC models

Here we propose four mixed-effects models: linear mixed-effects (LME), regularized mixed-effects (RME), multilevel mixed-effects (MME), and regularized multilevel mixed-effects (RMME). Each of the four models addresses the limitations and drawbacks we discussed earlier, and provides incremental improvements (Table 3).

**Table 3.**
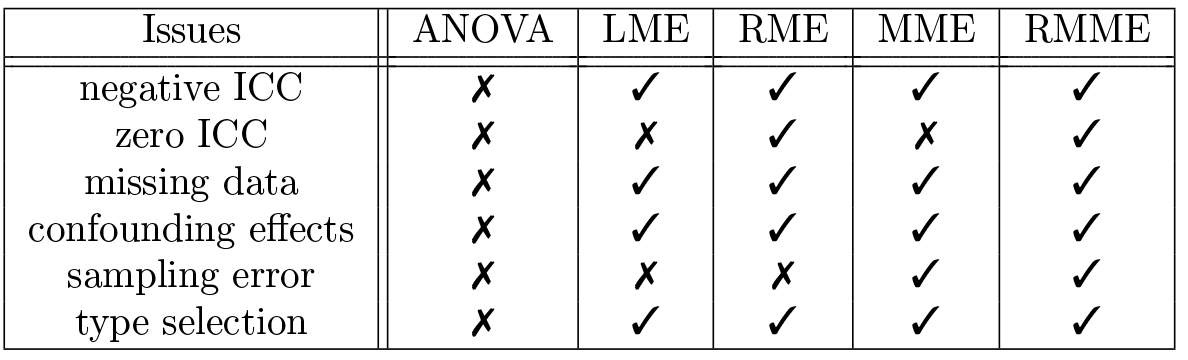
Issues in ICC computations under ANOVA, and summary of which mixed-effects models can (✓) or cannot (✗) address each.

| Issues | ANOVA | LME | RME | MME | RMME |
| --- | --- | --- | --- | --- | --- |
| negative ICC | ✗ | ✓ | ✓ | ✓ | ✓ |
| zero ICC | ✗ | ✗ | ✓ | ✗ | ✓ |
| missing data | ✗ | ✓ | ✓ | ✓ | ✓ |
| confounding effects | ✗ | ✓ | ✓ | ✓ | ✓ |
| sampling error | ✗ | ✗ | ✗ | ✓ | ✓ |
| type selection | ✗ | ✓ | ✓ | ✓ | ✓ |

### Linear mixed-effects (LME) modeling

Whenever multiple values (e.g., effect estimates from each of two scanning sessions) from each measuring entity (e.g., subject or family) are correlated (e.g., the levels of a within-subject or repeated-measures factor), the data can be formulated with an LME model, sometimes also referred to as a multilevel or hierarchical model. One natural extension to the ANOVA modeling in (10), (1), and (6) is to simply treat the model conceptually as LME, reformulating neither the equations nor their ICC definitions. This LME approach for ICC has previously been implemented in the program 3dLME (Chen et al., 2013) for voxel-wise data analysis in neuroimaging. For example, the ICC(2,1) model is an LME case with two crossed random-effects terms, whose applications can be seen under other circumstances such as inter-subject correlation analysis (Chen et al., 2016) and psycholinguistic studies (e.g., Baayen et al., 2008).

However, the application of LME methodology to ICC does not stop at the conceptual level, and in fact it has several advantages in some aspects of computation where limitations are present under the ANOVA framework. Specifically, the variances for the random effects components and the residuals are directly estimated through optimizing the restricted maximum likelihood (REML) function, and thus the ICC value is computed with variance estimates 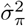,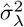 and 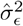, through the definitions in (11), (2), or (7), instead of with their counterparts with MS terms, (12), (3), or (8), under ANOVA. Therefore, in conjunction with the theoretical quantities, the estimated ICCs are nonnegative by definition, avoiding the interpretability difficulties that ANOVA-based estimates can present when negative. Similarly, the two F-statistic formulas, (5) and (9), can be expressed in terms of variance estimates as well,

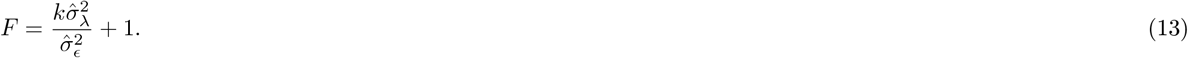

Another convenient byproduct from the LME model interpretation of (1) for ICC(2,1) is that the ICC for the absolute agreement between any two measuring entities (e.g., subjects) can be easily obtained through ratio among the variances,

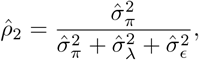

which is usually not discussed under the ANOVA framework.

In regard to the type selection choice between ICC(2,1) and ICC(3,1), which is an ambiguous one to some extent in the ANOVA framework, in the LME model (6) the fixed effects associated with the levels of the repeated-measures factor A can now be directly examined through statistical assessment: If the fixed effects *b*_*i*_ can be deemed negligible (i.e., no confounding effects nor systematic differences across the factor A levels), then ICC(2,1) offers a better metric because the model (1) is more parsimonious. We will elaborate this point later through an experimental dataset.

Furthermore, missing data can be naturally handled in LME because parameters are estimated through the optimization of the (restricted) maximum likelihood function, where a balanced structure is not required. As long as the missing data can be considered to occur randomly without structure (i.e., no systematic pattern exists among the missing data), and there are enough measuring entities present (e.g., *n* ≥ 10) for each level of the repeated-measures factor, then the computation can still be performed.

In addition, the extension to incorporate confounding effects is readily available through adding more fixed-effects terms into the model. For instance, the ICC(2,1) model (1) can be expanded to an LME model with two crossed random-effects components,

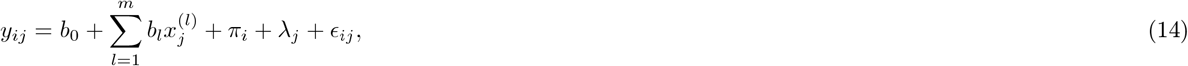

where *x*^(1)^,*x*^(2)^, …, *x*^(*m*)^ are *m* explanatory variables (e.g., sex, age) that can be either categorical or quantitative, and *b*_1_,*b*_2_, …,*b*_*m*_ are their corresponding fixed effects.

For the convenience of further discussion, we adopt the conventional LME platform for ICC with a more generic and inclusive formulation (Pinheiro and Bates, 2004; Chen et al., 2016) than (10), (1), or (6). The following formulation contains all of the aforementioned models as special cases, and will be further discussed hereafter:

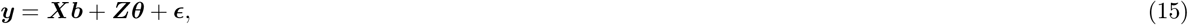

where the known vector *y*_*kn*×1_ = *vec*(*y*_*ij*_) is the vectorization or column-stacking of effect estimates *y*_*ij*_; ***b***_*m*+1_ contains the unknown fixed-effects to be estimated; the known model (or design) matrix ***X***_*kn*×(*m*+1)_ codes the explanatory variables for the fixed effects; the known model (or design) matrix ***Z*** is usually a column-wise subset of ***X***; *θ* contains the unknown random effects to be estimated; and *ϵ*_*kn*×1_ contains the unkown residuals. The distributional assumptions are that θ ~ *N*(0, ***V***),*andϵ* ~ *N*(0, ***R***), where ***V*** and ***R*** are the unknown variance-covariance matrices for the random effects *θ* and residuals *ϵ*, respectively; also, *θ* and *ϵ* are independent of each other (i.e., *Cov*(*θ*, *ϵ*) = 0). For the ICC context with no missing data, 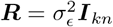 for ICC(1,1) and ICC(3,1), 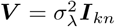 and for ICC(2,1), 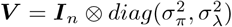.

For those ANOVA cases where a negative ICC value would occur due to the subtraction between two MS terms, LME avoids negativity by having a lower boundary at 0, via a positive definiteness for the variance-covariance matrix when estimating the variance components through optimizing the likelihood function within the nonnegative domain of the variance components. With these considerations in mind, the question for those ambiguous values in the LME framework becomes: are those regions in the brain fully uncorrelated across the levels of the repeated-measures factor; or is the zero variance estimate simply some artifact providing a zero; or is it a consequence of convergence failure in the optimization algorithms when solving LME? This is addressed by introducing an improvement to the conventional LME. We note that all of the advantages of LME over ANOVA discussed above also carry over to the other variants of mixed-effects models below.

## Regularized Mixed Effects (RME) model

When a zero variance estimate occurs, typically the corresponding likelihood profile is decreasing or close to a flat line at zero. Such a scenario may occur when the sample size is small or when the signal-to-noise ratio is low, derailing the LME capability to provide a meaningful variance estimate in this near-zero boundary value of zero. It is unfortunately not rare to have either a small number of subjects or a neuroimaging signal submerged with noise. Compared to a negative ICC value (or negative variance estimate 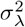), having a floor for 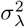 at 0 avoids an uninterpretable situation; however, even the zero estimate for ICC may be questionable to some extent: do we truly believe that all the subjects in a study have *exactly* the same BOLD response or average effect across the levels of factor A, as implied by such a value?

Variances in LME are estimated by optimizing REML, which is equivalent to the posterior marginal mode, averaging over a uniform prior on the fixed effects. To pull out of the trapping area surrounding the likelihood boundary, one possibility is to apply a weakly informative prior distribution for the variance parameters. This regularization approach can be conceptualized as forming a compromise between two competing forces: the anchoring influence of the prior information and the strength of the data. With a reasonable prior distribution, one may prevent a numerical boundary estimate from occurring by “nudging” the initial variance estimate by no more than one standard error, leading to negligible influence from the prior when the information directly from the data should be fully relied upon (Chung et al., 2013).

Here we adopt a weakly informative prior distribution, gamma density function (Chung et al., 2013), for a variance component v in the LME model (i.e., 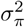 and 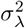 in (1), 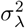 in (6)),

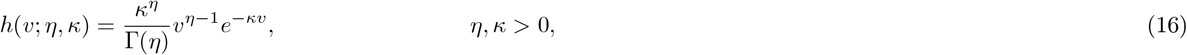

where *ŋ* and *k* are the shape and rate (inverse scale) parameters, respectively. In practice, the gamma density function has two desirable properties for our nudging purpose here: 1) a positive and constant derivative at zero when ŋ =2, guaranteeing a nudge toward the positive direction, and 2) *h*(0; *ŋ*, *k*) = 0 when *ŋ* > 1, allowing the variance to hit the boundary value of zero when the true value is zero. With *ŋ* = 2, the prior is uninformed (and improper) within (0,∞) when *K* approaches 0, and becomes gradually informative (but still weak) when *k* is away from 0 (Chung et al., 2013). Therefore, a parameter set of *ŋ* = 2 with a small rate parameter *k* produces a positive estimate for the variance but does not overrule the data itself.

Typical Bayesian methodology involves estimating the posterior distribution through sampling with simulations (e.g., Markov chain Monte Carlo). However, here the prior density can be directly incorporated into the likelihood function for LME because of the conjugacy of the exponential families, hence no simulations are required to nudge the variance estimate out of the boundary value for the ICC computation. As an added benefit, there is usually little extra computational cost added to the classic LME computations. In fact, an overall higher efficiency can be achieved in some circumstances, because the prior may speed up the convergence process otherwise stuck or slowed in the trapping area close to the boundary.

With both classic LME and its ‘nudging’ version, RME, there remains one last limitation mentioned in the Introduction to be overcome for ICC estimation: How can the LME model utilize the precision information (i.e., standard error) associated with the effect estimates from the individual subject analysis? Would the precision information provide a more accurate partitioning among the variance components including situations when ANOVA renders negative ICC or when LME forces 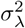 to be 0?

The effect estimates of interest in neuroimaging are usually not raw data or direct measures, but instead regression coefficients as output from a time series regression model; as such, these effect estimates have their own measurement uncertainties or confidence intervals. A close inspection of the LME model for ICC, (10), (1), or (6), reveals that the residual term *ϵ*_*ij*_ represents the uncertainty embedded in the effect estimates. An underlying assumption in the classic LME model holds that the measurement errors share the same uncertainty: 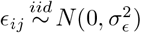. Since the effect estimates come from a time series regression model separately for each subject, their precision is not necessarily the same and may vary significantly across subjects for various reasons, including variations in trial sequence, data points, and outliers. Therefore, in practice, each effect estimate is associated with a different variance for its measurement error: 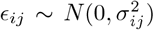. The true value of 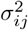 usually unknown, but its estimation is readily available, conveniently embedded in the denominator of the *t*-statistic for each effect estimate out of the individual subject analysis. Such an approach has previously been developed for simple group analysis, with the standard error for the effect estimate incorporated into the group model in neuroimaging (e.g., FLAME in FSL (Woolrich et al., 2004); 3dMEMA (Chen et al., 2012)).

## Multilevel mixed-effects (MME) modeling

Here we extend the LME model, (10), (1), or (6), through replacing the assumption 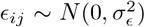 with an unknown parameter 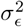 in the model by 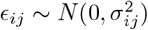, where the variance estimate 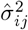 for 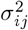 is assumed to be available, which is usually the case for task response. We call this approach multilevel mixed-effects (MME) modeling, with the term *multilevel* reflecting the fact that the modeling approach borrows part of a methodology typically adopted in robust meta analysis when summarizing across previous studies, each of which provided both effect estimate and its standard error. The MME counterpart for the standard LME formulation (15) is extended to have the assumption 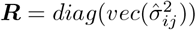.

Although computationally more sophisticated due to the involvement of more than one variance component in the case of the model (1) for ICC(2,1), the basic numerical scheme remains similar to our previous work for group analysis (Chen et al., 2012). That is, the variance components for the random effects are iteratively solved through optimizing REML, with the estimates 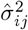 for measurement precision playing a weighting role: an effect estimate *y*_*ij*_ with a smaller (or larger) 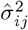 has a larger (or smaller) impact on the estimation of the variance components and fixed effects. It is this strategy of differential weighting that separates MME from the previous models in which each of the effect estimates is treated equally.

One adjustment for MME specific to the ICC case is the following. The variance for the residuals, 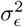 in the ICC definitions under all other models is no longer available, due to the replacement of the residual term with an unknown variance by the measurement error with an estimated variance. In its place, we substitute 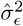 of with the weighted (or “typical”) average 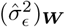 instead of the arithmetic average of the sample variances 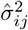 in light of the generic model (15) (Higgins and Thompson, 2002; Viechtbauer, 2010), where

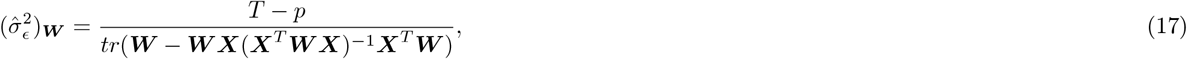
and *T* is the total number of data points in ***y*** (where *T* = *kn* if no missing data occur); *p* is the column rank of ***X***; *tr*() denotes the trace operator; ***X*** and 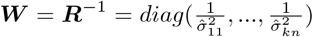 are the model matrix for the fixed effects in the model (15) and the weighting matrix, respectively. The differential weighting is reflected in the heterogeneous diagonals of the variance-covariance matrix ***R*** for the measurement errors, which becomes more transparent with a reduced form of 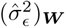 in the case of no missing data and no any explanatory variables, i.e., ***X*** = 1_*kn*×1_, in the model (15),

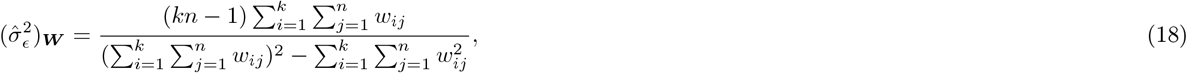

where the weights 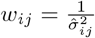.
*^a^ij*

## Regularized MME (RMME)

A zero variance estimate may still occur under MME for the same reason as in LME, namely that the corresponding likelihood profile peaks at zero. Similarly, the same gamma density function (16) can be adopted as a weakly informative prior distribution (Chung et al., 2013), for a variance component *v* in the MME model (e.g., see the example in (16) here), parallel to the extension from LME to RME. Here, as well, since no posterior samplings are involved in the process, the prior sometimes can even speed up the convergence process compared to MME. We note that RMME solves all the issues raised here, with the current ANOVA framework, as summarized in Table 3.

## Performance comparisons among the models

### Model implementations

Here, we have discussed three types of ICC, each of which can be estimated through five modeling strategies (ANOVA, LME, RME, MME, and RMME), leading to a total of 15 scenarios with an accompanying *F*-statistic defined by either (5) or (9). These 15 models and the corresponding *F*-statistics are all implemented in a publicly available program `3dICC` in AFNI, making use of the *R* packages *psych* (Revelle, 2016), *lme4* (Bates et al., 2015), *blme* (Chung et al., 2013), and *metafor* (Viechtbauer, 2010). The program `3dICC` takes a three-dimensional dataset as input for whole-brain voxel-wise analysis, but it can also be utilized to analyze a single voxel, region, or network. Additionally, parallel computing is available with multiple CPUs through the *R* package *snow* (Tierney et al., 2016). For each ICC model except ANOVA, the fixed effects (intercept or group average effect for each of the three models, as well as additional comparisons among factor A levels for ICC(3,1)) and their corresponding *t*-statistics are provided in the output from `3dICC`, in addition to ICC and the associated F-statistic value.

### Experimental testing dataset

To demonstrate the performances of our four proposed modeling approaches in comparison to ANOVA, we utilize experimental data from a previous FMRI study (Haller et al., 2017). Briefly, 25 healthy volunteers (mean age = 13.97 years, SD = 2.22 years, range = 10.04-17.51 years; 60% female) were asked to judge the gender of happy, fearful and angry face emotions. Each emotion was displayed at three intensities (50%,100%, 150%). A neutral condition, representing 0% intensity, was included for each face emotion (i.e., three neutral subsets were created, one for each face emotion). MRI images were acquired in a General Electric 3T scanner (Waukesha, WI, USA), and the participants completed two MRI scanning sessions approximately two-and-a-half months apart (mean = 75.12 days, SD = 15.12 days, range: 47-109 days). The FMRI echoplanar images (EPIs) were collected with the following scan parameters: flip angle = 50 degrees, echo time = 25 ms, repetition time = 2300 s, 47 slices, planar field of view = 240 × 240 mm^2^, acquisition voxel size = 2.5×2.5×3 mm^3^, and three runs with 182 volumes for each in a total acquisition time of 21 minutes. The parameters for the anatomical MPRAGE images were: flip angle = 7°, inversion time = 425 ms, and acquisition voxel size = 1 mm isotropic.

The EPI time series went through the following preprocessing steps in AFNI: de-spiking, slice timing and head motion corrections, affine alignment with anatomy, nonlinear alignment to a Talairach template TT_N27, spatial smoothing with a 5 mm full-width half-maximum kernel and scaling by the voxel-wise mean. Individual TRs and the immediately preceding volume were censored if: 1) the motion shift (defined as Euclidean norm of the derivative of the translation and rotation parameters) exceeded 1 mm between TRs; or 2) more than 10% of voxels were outliers. Only trials with accurate gender identification were included in the final analysis, but incorrect trials were also modeled as effects of no interest. Separate regressors were created for each of 13 event types (i.e., 30 trials for each face emotion at each intensity, neutral trials represented by three regressors of 30 trials each, and incorrect trials). Six head motion parameters and baseline drift using third order Legendre polynomials were included as additional regressors. The two sessions were analyzed separately, but the three runs with each session were concatenated and then entered into a time series regression model with a serial correlation model of ARMA(1,1) for the residuals.

The effect estimate in percent signal change, combined with the variance for the corresponding measurement errors, for the neutral condition associated with angry face-emotion from the individual subject analysis, was adopted for comparing the five ICC models: ANOVA, LME, RME, MME, and RMME. ICC(1,1) is not applicable in this case, but both ICC(2,1) and ICC(3,1) and their *F*-statistics were computed for each of the five models; the session effect and the corresponding *t*-statistic were also examined. The runtime was about one hour for each of the analyses with 16 CPUs on a Linux system (Fedora 14) with Intel^®^ Xeon^®^ X5650 at 2.67 GHz.

### Model comparisons

ANOVA renders a substantial number of voxels with negative ICC values (first column in Panel A, Fig. 1; first row and first column in Fig. 3), while LME provides virtually the same ICC estimates as ANOVA, with primary difference that those negative ICC estimates are replaced with 0 (uncolored voxels in the second column in Panel B, Fig. 1; scatterplot cells (1, 2) and (2, 1) in Fig. 3). It is worth noting that a significant proportion of voxels with negative or zero ICC from ANOVA or LME appear in gray matter. For RME, we tested four different priors by varying the rate parameter *K* at values of 0, 0.1, 0.3, and 0.5 (with *ŋ* fixed at 2); their differences in ICC estimates across the four *K* values are negligible. Furthermore, the computation cost is a decreasing function of *K* (substantially highest at *K* = 0). In light of these results, we set an empirical prior of gamma density (16) at ŋ = 2, *K* = 0.5 for neuroimaging data.

**Figure 1:**
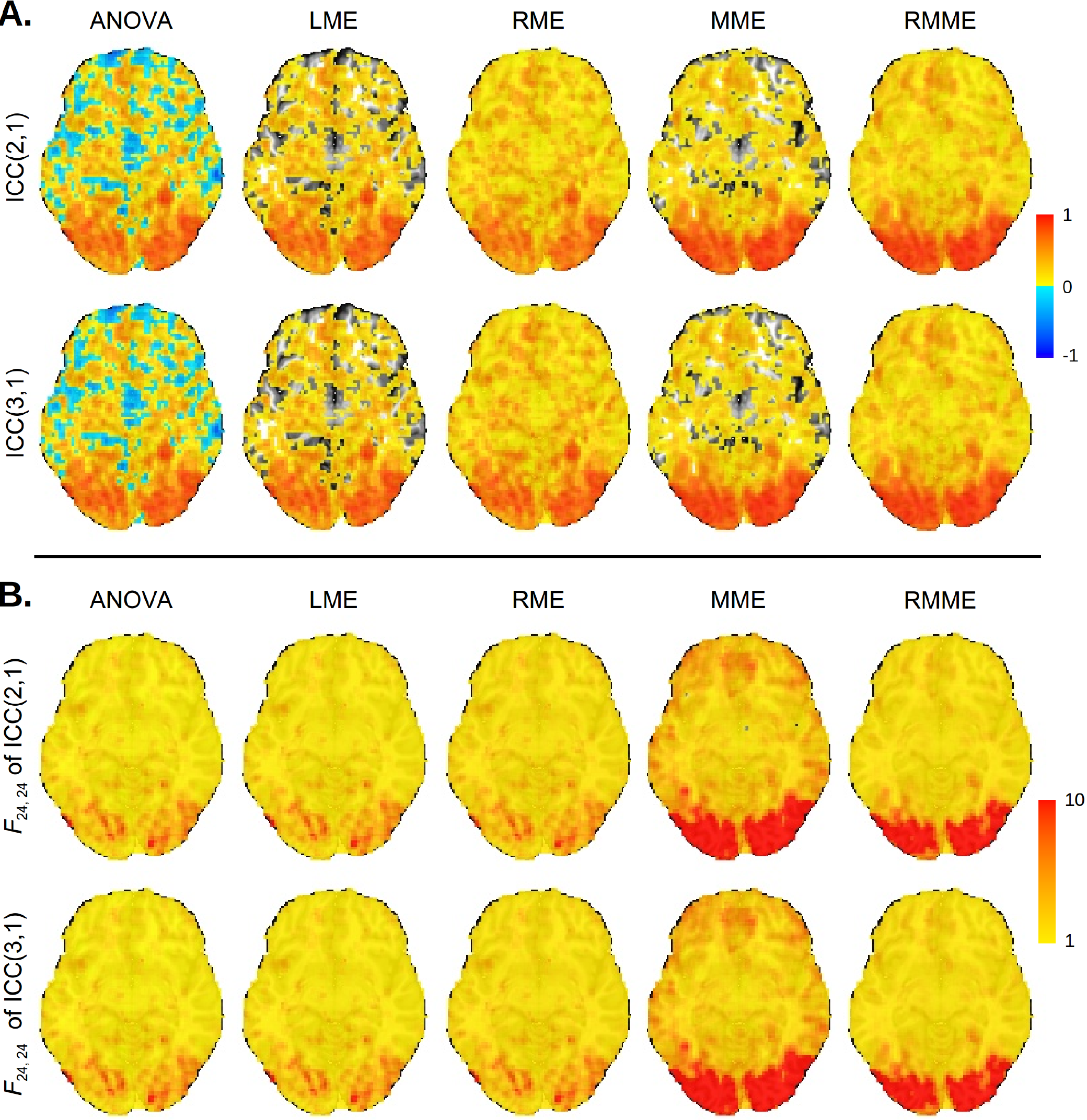
**Panel A.** ICC maps on an axial slice (*Z* = −6 mm, TT standard space; radiological convention: left is right) with a whole brain dataset through four models. The conventional ANOVA (first column) and LME (second column) rendered substantial number of voxels with negative (blue) and zero (not colored) ICC values, respectively. **Panel B.** The F-statistic maps corresponding to each of the ICC maps in Panel **A** are shown with colors in the range of [1, 10], with the lower bound of 1 defined by the formulation of *F*-statistic in (5). In each case, the degrees of freedom were (24, 24).

**Figure 2:**
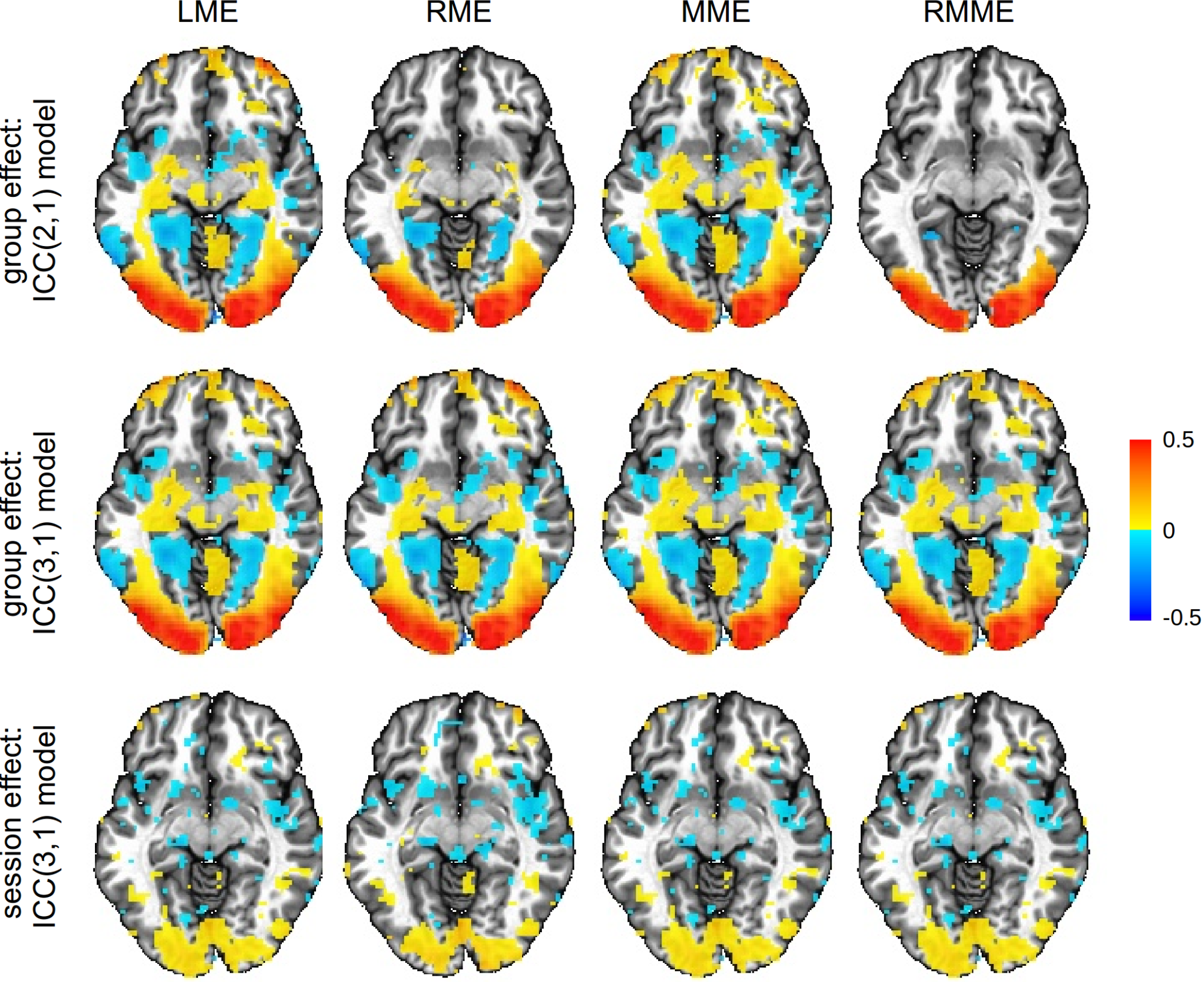
The first two rows show the group average effect on an axial slice (*Z* = −6 mm, TT standard space; radiological convention: left is right) with a whole brain dataset estimated through the four mixed-effects models. The third row is the session effect in the ICC(3,1) model. While the maps display the session effect, they are thresholded at a liberal significance level of 0.1 with 24 degrees of freedom for the corresponding *t*-statistic. The estimates for the fixed effects are not directly available from ANOVA, and therefore are not displayed. As the two regularization approaches estimate slightly higher variances under the ICC(2,1) model, some regions in the frontal area fail to survive the liberal thresholding for RME and RMME as shown in the first row. In contrast, all the four models demonstrate similar results under the ICC(3,1) model as shown in the second row.

**Figure 3:**
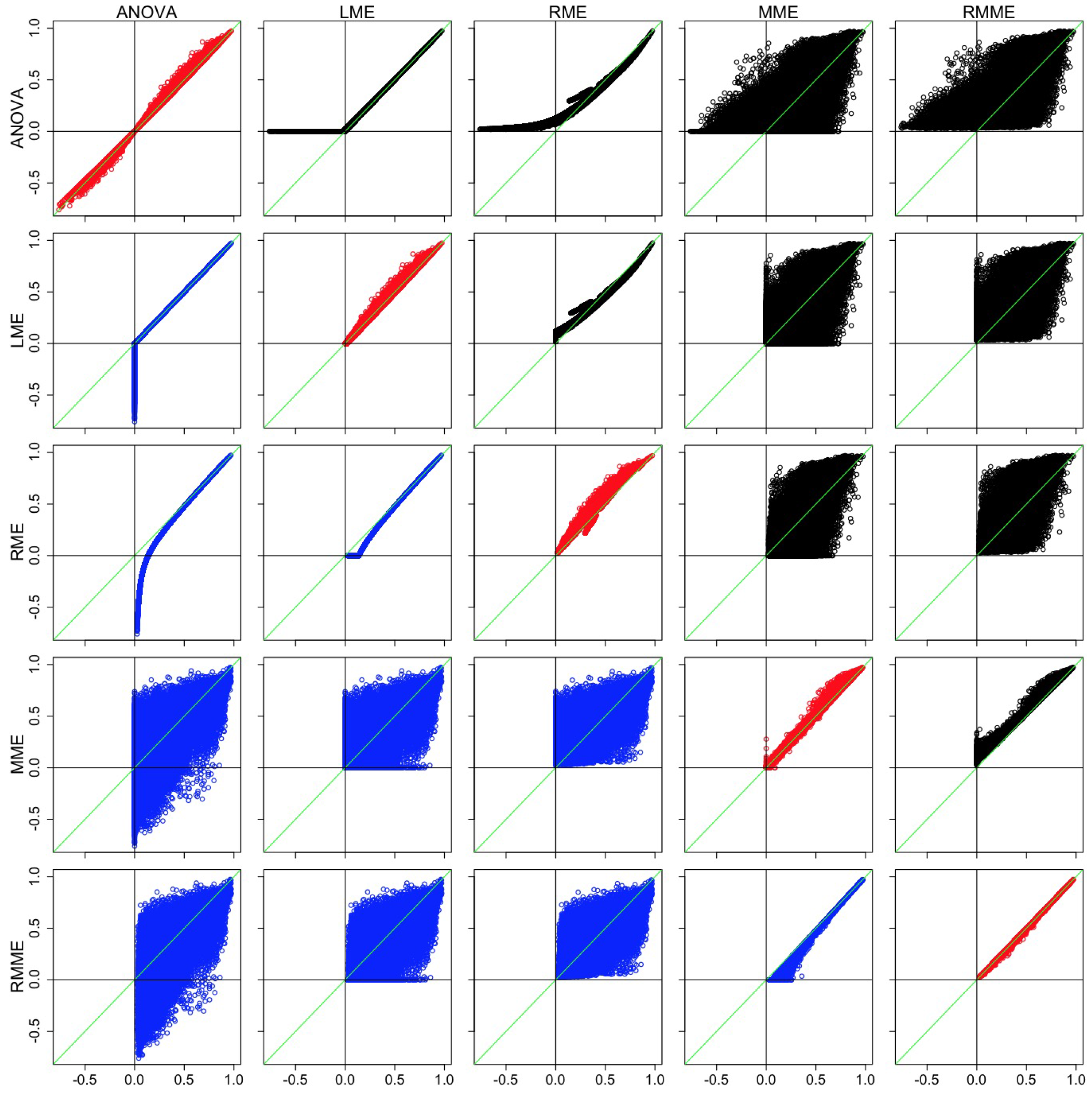
Comparisons across the five models and between ICC(2,1) and ICC(3,1) through scatterplots of ICC with the voxels in the brain. The *x* and *y* axes are a pair of ICC values between two of the five models with their labels shown at the left and top, respectively. The 10 combinatorial comparisons for ICC(2,1) are illustrated in the upper triangular cells in black; the 10 combinatorial comparisons for ICC(3,1) are listed in the lower triangular cells in blue; and the diagonals compare the two ICC types in red for each of the five models with ICC(2,1) and ICC(3,1) as *x* and *y* axes, respectively. In each plot the green diagonal line marks the situation when ICC(2,1) = ICC(3,1).

For the voxels with negative ICC values from ANOVA or with zero ICC values from LME, RME offers positive but generally small ICC estimates; the nudging effect of RME is relatively small when the ICC value from ANOVA/LME is positive but small (less than 0.3), and it is negligible when the ICC value from ANOVA/LME is moderate or large (larger than 0.3) (third column in Panel A, Fig. 1; scatterplot cells (1, 3) and (3,1) in Fig. 3). Some of the ICC estimates from MME are larger to varying extent than ANOVA/LME/RME, while some are smaller (fourth column in Panel A in Fig. 1; “fat blobs” showing wide variation in the scatterplot cells (1, 4), (2, 4), (3, 4), and their symmetric counterparts in Fig. 3); there are a higher number of voxels with larger ICC from MME relate to ANOVA/LME/RME than those with smaller ICC values (slightly redder voxels in fourth column than the first three columns, Panel A in Fig. 1; more dots with ICC greater than 0.75 on the MME side of the green diagonal line in the scatterplot cells (1, 4), (2, 4), (3, 4), and their symmetric counterparts in Fig. 3). However, there are still a large fraction of voxels with zero ICC estimates from MME (uncolored voxels in the fourth column, Panel A, Fig. 1), although less than from LME. Last, RMME shares similar properties with MME relative to ANOVA/LME/RME (fifth column, Panel A, Fig. 1; scatterplot cells (1, 5), (2, 5), (3, 5), and their symmetric counterparts Fig. 3). The differences between RMME and MME parallel those between RME and LME (fifth column, Panel A, Fig. 1; scatterplot cells (4, 5) and (5, 4), Fig. 3); that is, RMME provides small but positive ICC estimates for those voxels with zero ICC under MME, and the nudging effect is negligible when the effect estimate is relatively reliable. The F-values across the five models follow roughly the same patterns as the ICC values (Panel B in Fig. 1) because the F formulation is closely related to its ICC counterpart, and the high reliability measure in the brain shares the same regions with group average effect revealed from the four mixed-effects models (first two rows in Fig. 2).

The differences between ICC(2,1) and ICC(3,1) (Panel A in Fig. 1; diagonal cells in Fig. 3) are mostly small for each of the five models except for those regions where session effect is substantial (third row, Fig. 2). Because any systematic differences between the two sessions are accounted in the ICC(3,1), but not ICC(2,1), model, the former tends to render slightly higher ICC estimates. This is demonstrated in Fig. 3, by the fact that most voxels are above the green diagonal line in the diagonal cells of scatterplots. In addition, one noteworthy phenomenon is that RMME narrows the differences between the two ICC types, as represented in Fig. 3 by the thinner band in scatterplot cell (5, 5) relative to the other diagonal cells.

To gain better insights into the relative performances of the five models, we demonstrate three scenarios with three representative voxels with Table 4 illustrating the differences among the five models and with Fig. 4 schematically showing the heterogeneity and heteroscedasticity at those three voxels (their effect estimates and their variances are listed in Appendix A). At voxel *V*_i_ (from the right middle occipital gyrus), the ICC estimates from the ANOVA, LME and RME approaches are similar across all the five models. Both MME and RMME estimates are not much different because those effect estimates are roughly equally precise (first two columns in the right panel). The ICC estimate for the two types do not differ much either as the difference between the two sessions is small (~0.01% signal change, Table 4).

**Fig. 4:**
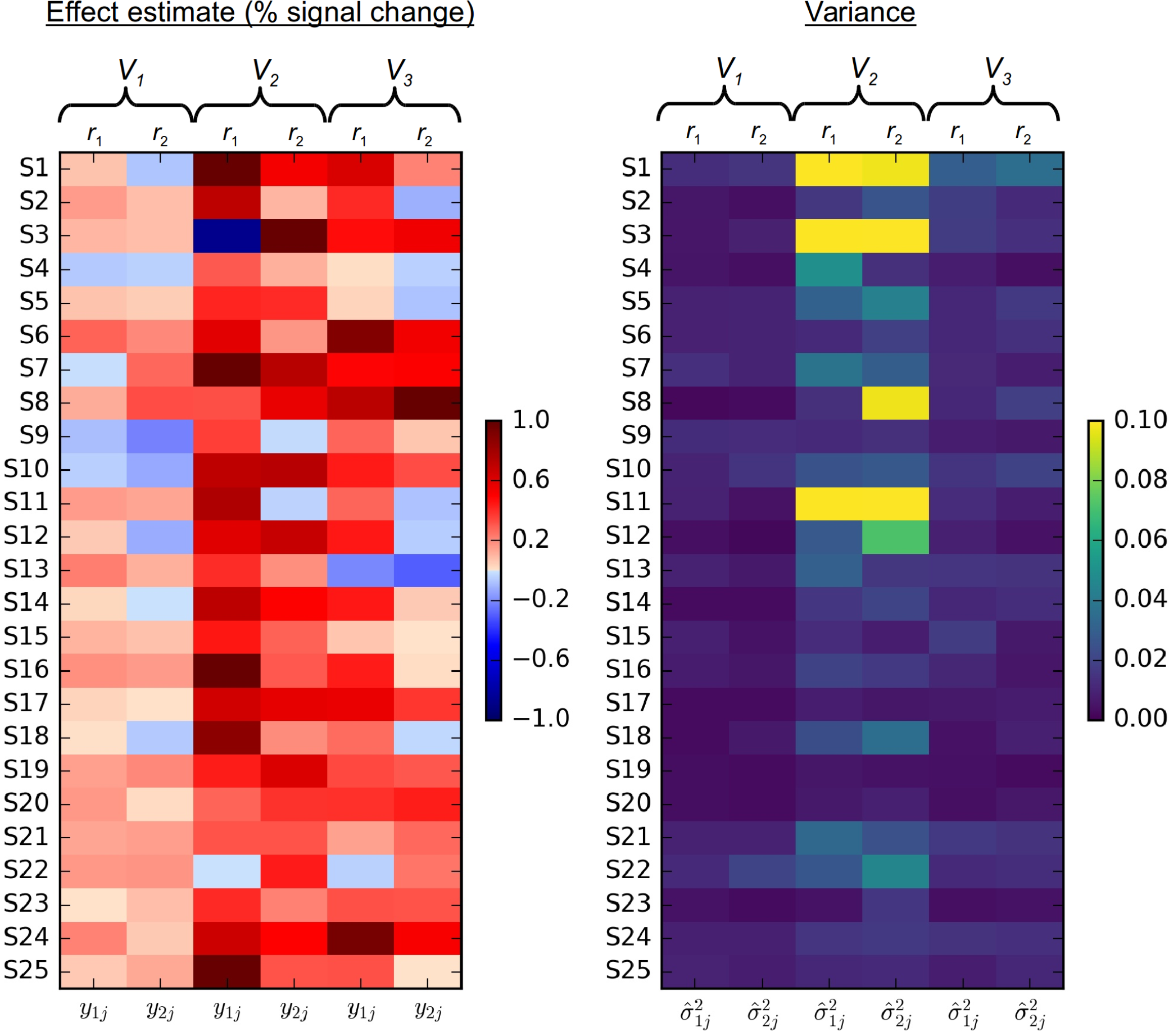
Example illustrations of heterogeneity and heteroscedasticity at three voxels with results shown in Table 4. Effect estimates (left) during the two sessions (*r*_1_ and *r*_2_) and the corresponding variances (right) for the measurement errors for the 25 subjects at three gray matter voxels (*V*_1_,*V*_2_, and *V*_3_) from the axial slice shown in Fig. 1. The real data are listed in Appendix A. Substantial session effect can be seen in voxel *V*_2_ (left matrix, middle two columns) while the session effect is negligible at voxels *V*_1_ and *V*_3_. A large amount of variability exists for the effect estimates at voxel *V*_1_ (left matrix, first two columns), leading to negative ICC estimate by ANOVA and zero by LME, while RME manages to deal with the degenerative situation with a small, but positive, ICC estimate. MME provides a positive and relatively large ICC estimate through weighting based on the precision information (right matrix, first two columns). The variability for the precision of the effect estimate is moderate at voxel *V*_2_ (right matrix, middle two columns), but minimal at voxel *V*_3_ (right matrix, right two columns).

**Table 4.**
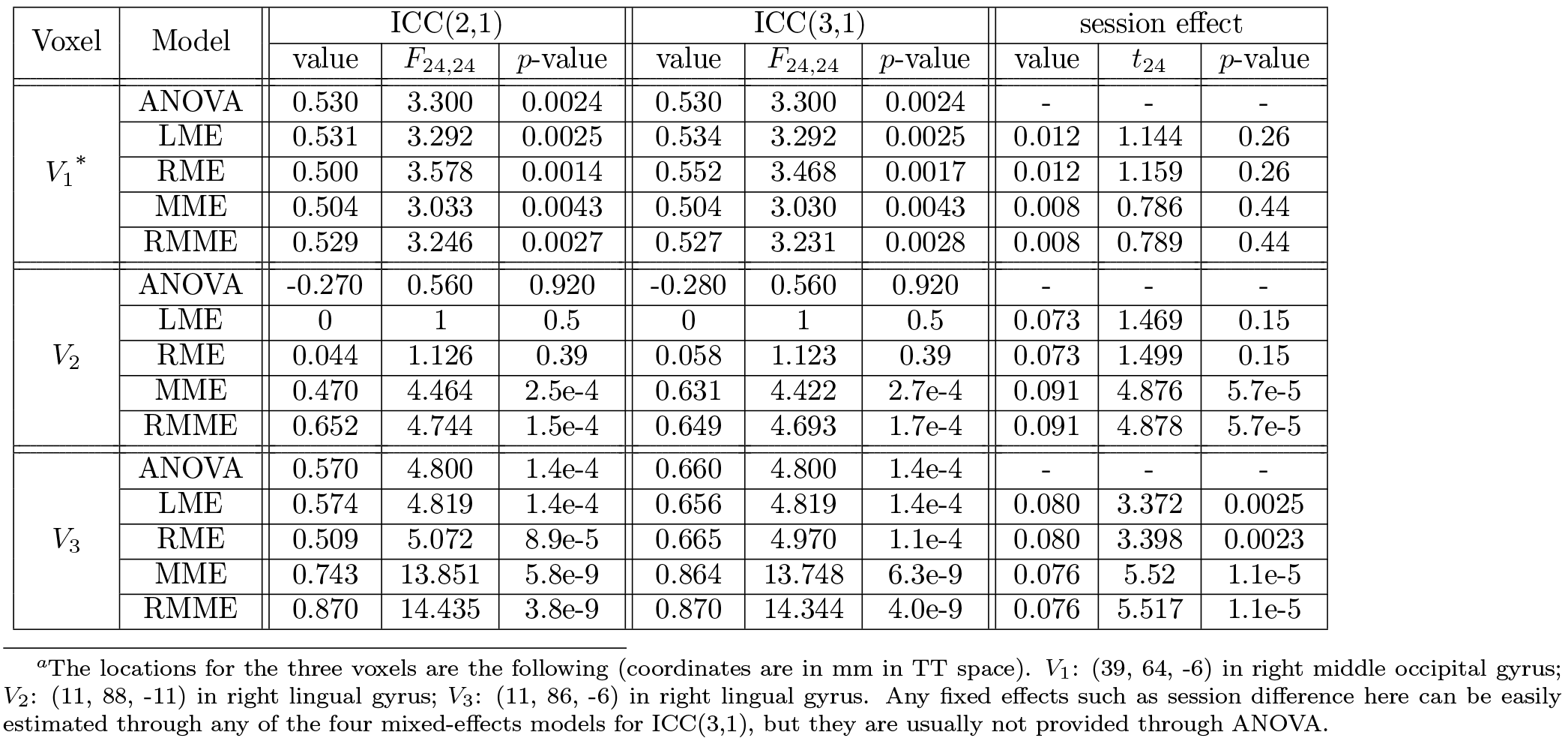
Results at three example voxels^a^ shown in Fig. 4

At voxel *V*_2_ (from the right lingual gyrus) ANOVA produces negative ICC values because MS_x_ is less than *M S*_*λ*_ in the corresponding ICC formulation (2) or (7). LME avoides negativity by taking the lower boundary value of zero that is allowed for variance, while RME renders a positive but small ICC estimate. In contrast, both MME and RMME achieve more accurate variance estimates in the sense that the availability of variances for measurement errors provides a way to reallocate or redistribute the components in the total variance. In other words, the substantial amount of heteroscedasticity as shown in Fig. 4 (third and fourth column in the right panel), allows differential weightings among the effect estimates with a higher emphasis on the more precise ones and downgrading for the less reliable ones. The dramatically different ICC values at voxel V2 from MME and RMME, relative to the three other ICC models, can be understood by examining the wide range of effect estimates as well as their heterogenous variances, as shown in the wide color spectrum of Fig. 4. It is worth noting that the presence of substantial session effect (close to 0.1%,*p* = 5.7 × 10^−5^) leads to a moderate difference between ICC(2,1) and ICC(3,1) for MME, but the analogous difference for RMME is negligible; that is, RMME tends to render similar ICC value between the two types regardless of the presence of session effect, just as shown in the whole brain data (scatterplot cell (5, 5) in Fig. 3).

Last, at voxel V3 (from the right lingual gyrus) ANOVA, LME and RME reveal moderately reliable effect estimates with similar ICC estimates. However, both MME and RMME render higher ICC values, due to the presence of moderate amount of heteroscedasticity, similar to the situation with voxel V2 even though less dramatic here (last two columns in both panels, Fig. 4). Also similar to voxel V_2_, the session effect (~0.08%, *p* =1.1 × 10^−5^) results in a lower estimate from MME for ICC(2,1) than ICC(3,1), but RMME estimates virtually the same reliability between the two ICC types. It is also interesting to note that, at both voxels V_2_ and V_3_, the *t*-statistic for the fixed effect of session is much higher when the precision information is considered than in the other two models, LME and RME (Table 4). This phenomenon demonstrates the potential impact and importance of including modeling precision in neuroimaging group analysis (Worsley et al., 2002; Woolrich et al., 2004; Chen et al., 2012).

## Discussion

Reliability is a crucial foundation for scientific investigation in general, and it has been a challenging issue and a hot topic for neuroimaging in particular over the years (e.g., Bennett and Miller, 2010). ICC offers a metric that can measure reliability under relatively strict circumstances. If the same scanning parameters are applied to the same cohort of subjects or families that undergo the same set of tasks, ICC shows the reliability across the levels of a categorical variable such as twins, sessions, or scanners. Nevertheless, in practice it is difficult to keep scanning circumstances perfectly constant across sessions in neuroimaging, thus systematic differences may slip in one way or another. Additionally, ICC can be applied to neuroimaging in the classic sense (i.e., inter-rater reliability or concordance), assessing the reliability between two diagnostic approaches (e.g., human versus automatic method) on a disease (e.g., schizophrenia, autism, or depression) or two different analytical methods applied to the same collection of data.

As shown here, there are three types of ICC, each of which can be estimated through various models such as ANOVA, LME, RME, MME, and RMME. On one hand, the ICC metric offers a unique approach to measuring the reliability of neuroimaging data under some well-controlled conditions; on the other hand, the investigator may still face a daunting job in deciding which ICC type, as well as which model, is most appropriate to apply in a given experiment or setup. We have presented the different formulations of each here, as well as demonstrated differences in data outcomes. We further discuss and summarize recommendations for model selections below.

### Considerations for effect estimates as inputs for ICC analysis

In practice, several factors can contribute to having poor data quality and accuracy in neuroimaging. Having relatively low signal-to-noise ratio is a major issue, and suboptimal modeling due to poor understanding of the major components in the signal is another. By some estimates, less than half (and in some cases down to 20-30%) of data variability can be accounted for in a typical FMRI data analysis at the individual level (Gonzalez-Castillo et al., 2017). Some rigor and standardization steps are required to achieve more accurate reproducibility and reliability. At present, the cumulative impact of altering preprocessing and modeling strategy in an analysis pipeline is largely unknown.

For example, it is well known that the absolute values of the FMRI-BOLD signal from the scanner have arbitrary units with scaling fluctuations among subjects, and therefore some kind of calibration is needed during preprocessing if the effect estimates are to be used in further analyses at the group level, for both typical group analysis as well as ICC estimation. Such a calibration should take into consideration the fact that the baseline varies across brain regions, subjects, groups, scanners and sites. However, radically different scaling strategies are in use, and their effects at the group level remain unclear. For instance, the global or grand mean scaling, as currently typically practiced and adopted by some software packages, can only account for part of the cross-subject variability; however, such approaches do not address the cross-region variability within an individual brain as a practical reality in neuroimaging scanning, and therefore either may lead to difficulty in interpreting and comparing the effect estimates. Analyses on networks including ICC and causal modeling would be affected as well. In contrast, voxel-wise scaling as a calibrator, even though imperfect due to the ambiguity in the baseline definition (Chen et al., 2017), provides a more accurate characterization than the alternatives. Because of these considerations, in general we recommend voxel-wise mean scaling during preprocessing, so that a directly interpretable and compatible measure in percent signal change can be adopted for the effect estimates that are to be taken for group analysis including ICC.

Lastly, each effect estimate in neuroimaging is typically derived from a model coefficient, and is intrinsically associated with some extent of uncertainty that may vary across subjects. As the precision information of the single subject effect estimate (embedded in the denominator of *t*-statistic) is required for the preferred MME and RMME, it is important to model the temporal correlation in the residuals at the individual level to avoid inflating the precision (or underestimating the uncertainty) for the effect estimate to be used in the ICC model.

### Which ICC model to use?

Among the thousands of voxels in a typical whole-brain neuroimaging dataset, negative ICC values unavoidably, and even frequently, show up, though they are usually not reported in the literature. The standard ICC values reported in most literature to date contains several aspects of ambiguity, greatly hindering meaningful interpretation. In addition, even when the ICC type is reported, its reason for selection, for example, between ICC(2,1) and ICC(3,1), is usually not clearly explained. In some cases, the chosen method is ill-suited to the given analysis scenario (e.g., if ICC(1,1) was used between two sessions).

When precision information for the measurement errors of the effect estimate is available, we recommend using RMME for the following three reasons: 1) the precision information offers a more robust estimate for ICC, as well as for the fixed effects in the model; 2) the regularization aspect of the approach leads to the avoidance of the uninterpretable situation of a negative ICC from ANOVA or an unrealistic zero ICC estimate from plain LME; 3) as demonstrated here (scatterplot cell (5, 5) in Fig. 3 and Table 4), RMME tends to be less sensitive to ICC type selection, rendering roughly the same ICC estimate regardless of the type the investigator adopts. When the precision information is unavailable (e.g., when one has the correlation value as the effect of interest from seed-based analysis or psycho-physiological interaction analysis), we recommend using RME because of its capability to provide a realistic ICC estimate when ANOVA (or LME) renders negative (or zero) ICC.

A general linear model (GLM), as extension to ANOVA, can accomodate between-subjects variables and assist the investigator in ICC type selection. Therefore, the GLM is an intermediate approach between ANOVA and LME. However, it still shares the other limitations with ANOVA in the following aspects: rendering negative or zero ICC, and being unable to handle missing data or to incorporate sampling errors.

The use of a regularization approach may raise questions regarding its insertion of arbitrariness into the estimation process with a prior for the variance components. Each variance component in the ICC formulation is estimated as a point value; however, it is worth noting that the value of a variance component usually does not exactly fall at its numerical estimate, but varies within some range (e.g., 95% central or uncertainty interval). The reason for a negative or zero estimate for a variance component lies in the methodology of replacing the variance by a point estimate; in other words, the substitution with a point estimate ignores the fact that there is uncertainty associated with the estimate. As the point estimate usually tends to be imprecise and underestimated, the negative or zero variance estimate or ICC should not be taken at the face value (Chung et al., 2013). A zero estimate for variance and ICC can also cause inflated inferences on the fixed effects such as group average as well as systematic differences across the factor A levels in the ICC(3,1) model. More fundamentally, simply forcing a negative ICC or the variance estimate for 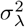 in (2) or (7) to zero leads to an unjustifiable claim that the effects from all the subjects are absolutely the same. On the other hand, the regularization approach can be conceptualized as a tug of war between the prior and data. As the results from our experimental dataset demonstrate here (e.g., Fig. 1), a weakly informative prior for ICC estimation can pull a degenerate situation (e.g., zero variance estimate) out of the boundary and render a more reasonable estimate, while little impact will incur when the data contain strong information, which would overrule the prior.

Four aspects of MME and RMME are worth highlighting here. First, these two multilevel approaches estimate ICC differently from the other three methods to some extent, as indicated in those “fat blobs” of Fig. 3 among cells (1,4), (2,4), (3,4), (1,5), (2,5), and (3,5) for ICC(2,1), and (4,1), (4,2), (4,3), (5,1), (5,2), and (5,3) for ICC(3,1). It should be stressed that the focus here is not on which method leads to a higher or lower estimate (nor should it be), but on which model provides a more accurate characterization about the reliability measure. Second, just as the levels of a within-subject factor are treated as simultaneous variables of a multivariate model (as in the AFNI group analysis program `3dMVM`) for repeated-measures ANOVA in neuroimaging group analysis (Chen et al., 2014), so are the factor A levels in MME and RMME for ICC(3,1) modeled here in a multivariate fashion with the flexibility in variance decomposition (Viechtbauer, 2010) and the capability to incorporate quantitative covariates in the presence of a repeated-measures factor. Third, the measurement errors associated with the factor A levels in a generic model of the form in (1) or (6) can be correlated when different tasks are intertwined in the experiment, thus the variance-covariance matrix ***R*** should be semi-definite in general. It is because of the presence of covariances in ***R*** that the modeling approach adopted in 3dMEMA of AFNI and FLAME of FSL cannot instead be utilized to perform a paired test in cases where the two effect estimates are entered separately as input (even if the program permits such an option), as the measurement errors corresponding to the two effect estimates are usually correlated. However, ***R*** is actually a diagonal matrix in the neuroimaging ICC context, because the measurement errors can be reasonably assumed to be well-approximated as independent with each other (e.g., across runs or sessions). Finally, nonparametric methods (e.g., bootstrapping, permutations) may offer a robust venue for controlling family-wise error for relatively simple models at the group level; however, under some scenarios parametric approaches provide more specific and accurate characterization about the effect of interest through some quantifying parameters (e.g., variance components in the ICC context) in the model, which are currently both valid and irreplaceable, as shown here with ICC computations through ANOVA, LME, RME, MME, and RMME, as well as the group analysis approach through incorporating the precision information (Worsley et al., 2002; Woolrich et al., 2004; Chen et al., 2012).

### Which ICC type to adopt?

As there is usually one single effect estimate for each subject per scanning situation, our discussion here focuses on single-measurement ICC. Among the three ICC types, ICC(1,1) is likely the easiest to list the scenarios in which it can be applied. Primarily, it can be used when there is no apparent distinction for a sequence of the levels for the factor that is associated with the multiple measurements in the ICC model. It typically applies to the situation of studying twins, for example.

In contrast, the other two types are utilized for scenarios such as having two or more runs, sessions, scanners, sites, or between parent and child. The reliability from ICC(2,1) represents “absolute agreement,” to the extent that the values exactly match between any two levels of the factor A, while ICC(3,1) shows the consistency or the extent that the effect estimates match across the factor A levels *after* accounting for potential systematic differences or other fixed effects. In other words, if the systematic differences across the levels of factor A are negligible, then the two ICC estimates would be similar. On the other hand, if the systematic differences or confounding effects are substantial, then the ICC values tend to diverge to some extent, and they lead to different interpretations. However, the existence of systematic differences itself warrants further exploration about the source or nature of those fixed effects (e.g., habituation or attenuation). For example, what is the association between the ICC map and the activation map (i.e., intercept in the model (14) or (15))?

Due to the simultaneity of analyzing all the voxels, it is unrealistic to choose one ICC type for some regions while selecting another for the rest. Per the discussion here between ICC(2,1) and ICC(3,1), we generally recommend the latter for whole brain analysis. In doing so, potential fixed effects are properly accounted for. More importantly, it is not the ICC interpretation in the sense of absolute agreement that is generally of primary importance, but the extent of agreement after potential fixed effects are all explained. Furthermore, with ICC(3,1), the investigator can directly address the following questions: 1) which brain regions show systematic effects across the levels of factor A? 2) how do those systematic effects correspond to the ICC maps in these regions? and 3) are the systematic effects related to some confounding effects such as habituation or attenuation?

## Result reporting

Clear and accurate scientific reporting is important for result reproducibility, reliability and validity as well as for meta-analysis. The present reporting conventions in neuroimaging are especially discouraging, with a lopsided focus on statistics alone, e.g., due to oversight or limitations in software implementations (Chen et al., 2017b), leading to incomplete reporting throught the literature. It cannot be emphasized enough that the effect estimates involved in a study should be reported in addition to the statistical significance, and the same emphasis should be applied to ICC analysis. Specifically, the investigator should explicitly state the ICC type and the model adopted, as well as the justification for such choices. One may notice that the ICC formulation for ICC(1,1) in (11) and (12) is exactly the same as ICC(3,1) in (7) and (8), which means that reporting the ICC formula would not be enough to reveal the whole picture because the two underlying models, (10) and (6), are dramatically different (and so are the two resulting ICC estimates).

With regards to the criteria for reliability, a loose rule of thumb has been suggested for ICC values as the following (Cicchetti, 1994): [0, 0.4), poor; [0.4, 0.6), fair; [0.6, 0.75), good; and [0.75, 1], strong. One cautionary note is that a low ICC does not always mean poor reliability: it is possible that some confounding effects are not accounted for in the model. For the statistical significance of ICC, one may use the Fisher transformation (4), or preferably, the F-statistic (5) and (9). Nevertheless, the F-statistic is not necessarily a basis for clusterization, but together with the ICC value, it serves as some auxiliary information to gauge the reliability about the effect of interest from the conventional analytical pipeline. Finally, all the fixed effects including the intercept (group average) are crucial part of the model and should be discussed and explained in the paper as well, as exemplified here in Fig. 1 and Fig. 2.

## Conclusion

One potential problem with the classic definition of ICC is that negative ICC values may appear, a scenario which is almost certain to show up in a whole brain neuroimaging analysis. Here we extend the conventional ICC definition and its computations to the frameworks of LME, RME, MME, and RMME modeling to address this issue and other difficulties. Such an extension not only offers wider modeling flexibility such as the inclusion of various fixed effects, but also avoids the interpretability problem of negative ICC values under ANOVA. We offer our recommendations in model adoption and ICC type selection as well as in result reporting. All modeling strategies and ICC types as well as the estimation and statistic testing for the fixed effects are currently available in the AFNI program `3dICC`.

## Acknowledgments

The research and writing of the paper were supported by the NIMH and NINDS Intramural Research Programs (ZICMH002888) of the NIH/HHS, USA. We are indebted to Wolfgang Viechtbauer and Vincent Dorie for their precious help and for the R packages *metafor* and *brms*, which they build and maintain, respectively.

## Appendix A. FMRI processing

The general sequence of FMRI data preprocessing steps was described in the subsection **Experimental testing dataset** under the section **Performance comparisons among the models**. However, for greater specificity and reproducibility, in this Appendix we also include the exact *afni_proc.py* command in AFNI (version AFNI_16.3.06) that was implemented to create the full processing pipeline. While there are several processing steps specified, each with many user-chosen options, it is possible to provide the exact pipeline in a succinct manner because the processing steps and options were created and specified using *afni_proc.py* in AFNI. This tool permits the user full freedom to tailor a desired pipeline that may be reliably duplicated for the entire group, stored for future reference and published with a study for unambiguous description.

## Appendix B. Data at three voxels from 25 subjects with two sessions that are illustrated in Fig. 4 and Table 4

**Table 5:**
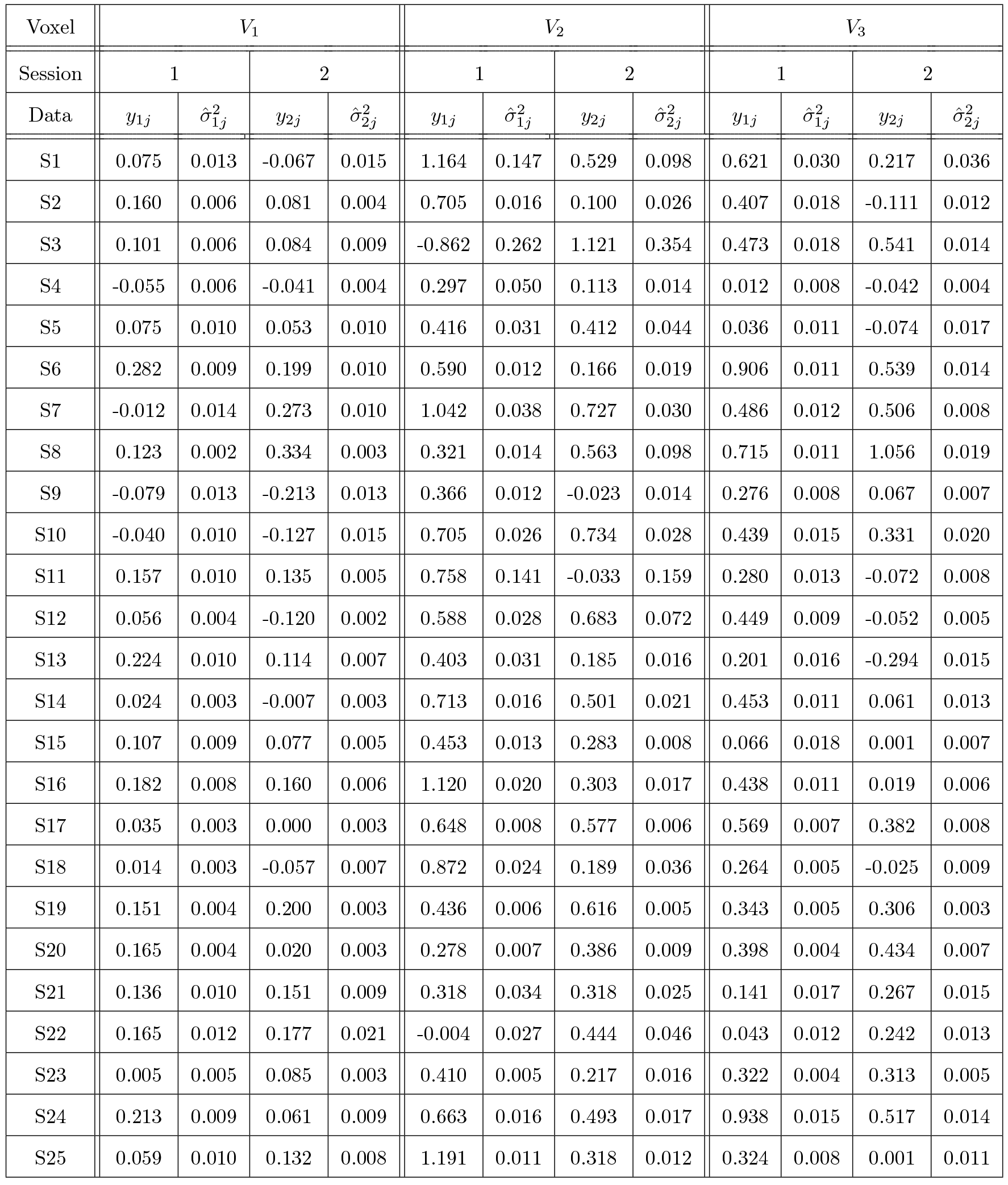
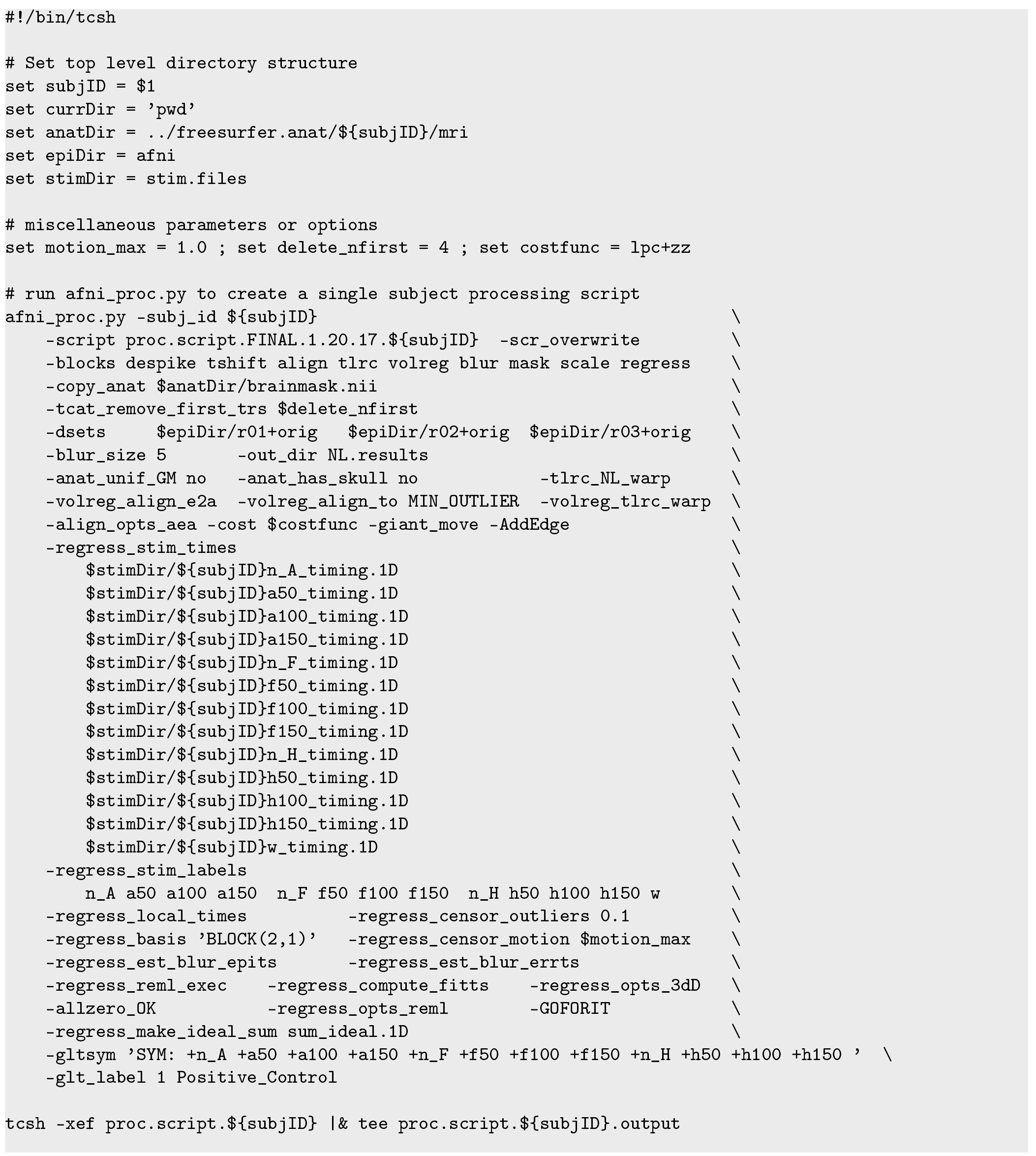
A compact tcsh script that contains the succinct, selected *afni_proc.py* command used to generate the full processing pipeline (>500 lines) in AFNI for this study. To implement across the group, one simply loops through a list of subjects, entering the given file name as the sole command line argument, which is passed to the variable $subjID. Here, stimulus variables are encoded as: h = happy, n = neutral, f = fearful, w = wrong; and each is followed by the duration (50, 100, 150) s.

## Footnotes

1 For simplicity, the notations for the model terms and for the corresponding variance and MS terms in the ICC formulas are undifferentiated across models. To avoid confusion, we emphasize that the values for the same notation term may change across models. For example, residuals *ϵ*_*ij*_ and its variance estimate 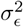. may differ, depending on how the effects associated with factor A are modeled.

2 One exception is that, when a subject-grouping factor (e.g., males versus females) is considered, it is possible to construct the involved MS terms in the special case of having exactly equal number of subjects across all groups. However, even for such a balanced scenario, specific MS terms would have to be derived for each ICC computation formula.

3 Precision is defined as the reciprocal of the variance.

4 Without loss of generality, the dimensions shown here for vectors and matrices are assumed to have no missing data, by default. When missing data occur, the dimensions can be adjusted accordingly.

5 The *F*-statistic is not exact in the cases of RME, MME, and RMME. However, it is important to note that the *F*-statistic would be an approximation too even for ICC(1,1) and ICC(3,1) under ANOVA and LME since the measurement errors are ignored.

6 Despite the concatenation, the discontinuities across runs were properly handled (Chen et al., 2012).

7 It might be neurologically more interesting to show the ICC in a few regions, but here we chose these three voxels instead of regions to demonstrate their subtle differences across ICC types as well as models.

8 Due to the complexity of handling within-subject factors, some group analyses in published Neuroimaging literature are still not performed correctly, leading to substantial number of publications with inflated statistical inferences (McLaren, 2010; Chen et al., 2014).

9 Instead, such a paired test can be validly performed by taking the contrast and its standard error as input.

10 To save space, the data shown here are rounded to the nearest thousandth, therefore there may have some small differences with the results listed in Table 4.

